# Evaluating intraspecific genetic diversity of a fish population using environmental DNA: An approach to distinguish true haplotypes from erroneous sequences

**DOI:** 10.1101/429993

**Authors:** Satsuki Tsuji, Masaki Miya, Masayuki Ushio, Hirotoshi Sato, Toshifumi Minamoto, Hiroki Yamanaka

## Abstract

Recent advances in environmental DNA (eDNA) analysis using high-throughput sequencing (HTS) provide a non-invasive way to evaluate the intraspecific genetic diversity of aquatic macroorganisms. However, erroneous sequences present in HTS data can result in false positive haplotypes; therefore, reliable strategies are necessary to eliminate such erroneous sequences when evaluating intraspecific genetic diversity using eDNA metabarcoding. In this study, we propose an approach combining denoising using amplicon sequence variant (ASV) method and the removal of haplotypes with low detection rates. A mixture of rearing water of Ayu *(Plecoglossus altivelis altivelis)* was used as an eDNA sample. In total, nine haplotypes of Ayu mitochondrial D-loop region were contained in the sample and amplified by two-step tailed PCR. The 15 PCR replicates indexed different tags were prepared from the eDNA sample to compare the detection rates between true haplotypes and false positive haplotypes. All PCR replications were sequenced by HTS, and the total number of detected true haplotypes and false positive haplotypes were compared with and without denoising using the two types of ASV methods, Divisive Amplicon Denoising Algorithm 2 (DADA2) and UNOISE3. The use of both ASV methods considerably reduced the number of false positive haplotypes. Moreover, all true haplotypes were detected in all 15 PCR, whereas false positive haplotypes had detection rates varying from 1/15 to 15/15. Thus, by removing haplotypes with lower detection rates than 15/15, the number of false positive haplotypes were further reduced. The approach proposed in this study successfully eliminated most of false positive haplotypes in the HTS data obtained from eDNA samples, which allowed us to improve the detection accuracy for evaluating intraspecific genetic diversity using eDNA analysis.

## Introduction

Genetic diversity is a key component of biodiversity and is necessary for species’ adaptation to changing natural and human-induced selective pressures (Allendorf et al. 2012; Laikre et al. 2016). Intraspecific genetic diversity of a fish population has typically been analyzed by capturing individuals using traditional methods, such as baited traps, casting nets, and electrofishing, followed by genetic analysis of each individual. Using restriction-site Associated DNA Sequencing (RAD-seq) and Multiplexed ISSR Genotyping by sequencing (MIG-seq) the traditional sampling approaches allow us to determine more detailed insights into population genetic diversity comparted to haplotypes (cf. Baird et al. 2008, Suyama et al. 2015). However, these methods have at least two major limitations: 1) use of traditional sampling approaches causes damage to the target organisms, and 2) large sampling efforts are necessary to provide an accurate estimate of intraspecific genetic diversity across an entire population. Traditional capture methods may threaten the persistence of species or population, particularly for rare and endangered species. In addition, insufficient sampling might lead to an underestimation of intraspecific genetic diversity of a population (Xing et al. 2013). These limitations may reduce the feasibility of surveys and increase the uncertainty of results.

Environmental DNA (eDNA) is DNA released from organisms into the environment (e.g. soil, water, and air), and it originates from various sources, such as metabolic waste, damaged tissues or sloughed skin cells (Kelly et al. 2014). Environmental DNA analysis has recently been used to detect the distribution of macroorganisms, particularly those living in aquatic habitats (Ficetola et al. 2008; Lodge et al. 2012; Thomsen et al. 2015). Environmental DNA analysis allows for non-invasive and cost-effective detection of the presence of a species in a habitat because only collection of water samples is required in the field instead of capturing and/or observing the target species (Thomsen et al. 2015). Because of this advantage and tis high sensitivity, eDNA analysis has frequently been applied for the detection of not only common species but also rare and endangered species (Fukumoto et al. 2015; Ishige et al. 2017; Katano et al. 2017; Rees et al. 2014; Thomsen et al. 2012). In addition, an approach involving eDNA metabarcoding using high-throughput sequencing (HTS) can effectively and comprehensively reveal the aquatic community structure, and thus, it has been gaining attention as a powerful tool for biodiversity monitoring (Kelly et al. 2014; Miya et al. 2015; Thomsen et al. 2012; Yamamoto et al. 2017).

Current applications of eDNA analysis have been limited mostly to the detection and identification of species (e.g. Rees et al. 2015; Thomsen et al. 2015). However, eDNA analysis can potentially be extended to the evaluation of intraspecific genetic diversity, because eDNA released from multiple individuals coexist in a water sample. Recently, Uchii et al. (2016 and 2017) have developed a method using cycling probe technology and real-time PCR to quantify the relative proportion of two different genotypes of common carp *(Cyprinus carpio)* based on a single nucleotide polymorphism (SNP). These studies revealed that the SNP genotypes were present in the sampled water. Furthermore, Sigsgaard et al. (2016) and Parsons et al. (2018) applied eDNA analysis for estimating of intraspecific genetic diversity in whale shark *(Rhincodon typus)* and harbour porpoise *(Phocoena phocoena)* populations, respectively, and found multiple haplotypes that had been identified previously from tissue-derived DNA by Sanger sequencing. These findings show the effectiveness of eDNA analysis for analyzing intraspecific genetic diversity of target species. However, caution should be exercised during the use of HTS for intraspecific genetic diversity, because HTS data usually include many erroneous sequences that are generated during PCR and sequencing (Coissac et al. 2012; Edgar et al. 2016a; Schloss et al. 2011).

Researchers have tried to address the issue of erroneous sequences using multiple approaches, containing the use of high-fidelity DNA polymerase in PCR, quality filtering based on base-call scores and/or clustering of sequences into operational taxonomic units (OTUs, OTU methods). The use of high-fidelity DNA polymerase in PCR contributes to decreased errors in PCR products (Ramachandran et al. 2011), but it does not completely prevent the errors. The OTU methods involve clustering of sequences that are less different from each other than a fixed similarity threshold (typically 97%; Callahan et al. 2016; Hughes et al. 2017). Thus, true haplotypes that are similar to each other are clustered into a single OTU, leading to incorrect evaluations of intraspecific genetic diversity. Therefore, OTU methods cannot be applied to analyze intraspecific genetic diversity. To evaluate intraspecific genetic diversity using eDNA samples, it is necessary to develop effective novel approaches to eliminate erroneous sequences inherent to HTS.

Intraspecific genetic diversity of a population might be analyzed more effectively by the use of amplicon sequence variant (ASV) methods, which have recently been developed in the fields of microbiology for correcting erroneous sequences derived from HTS data (e.g. Callahan et al. 2017 and references therein). ASV methods resolve amplicon sequence variants (ASVs) in the sample without imposing the arbitrary dissimilarity thresholds that define OTUs. As a core process of an ASV method, ‘denoising’ is performed using a de novo process in which biological sequences are distinguished from errors on the basis of, in part, the expectation that biological sequences are more likely to be repeatedly observed than erroneous sequences (e.g., DADA2; Callahan et al. 2016 and UNOISE2/3 by Edgar et al. 2016a). The sensitivity and accuracy of ASV methods with respect to correcting erroneous sequences have been shown to be better than those of OTU methods (Callahan et al. 2016; Edgar et al. 2016; Eren et al. 2013; Eren et al. 2015; Needham et the al. 2017). The high resolution of biological sequences afforded by ASV methods has the potential to improve the accuracy of evaluating intraspecific genetic diversity inferred from eDNA.

The purpose of this study was to propose an approach for eliminating false positive haplotypes derived from erroneous sequences in HTS data obtained from an eDNA sample and to demonstrate the usefulness of eDNA analysis for the evaluation of intraspecific genetic diversity of a fish population. In this study, we examined genetic diversity in the Ayu *(Plecoglossus altivelis altivelis)* fish, an important fishery target in Japanese inland waters whose genetic diversity has been evaluated in previous studies (e.g. Iguchi et al. 2002, Takeshima et al. 2016). First, we examined whether we could correctly determine the mitochondrial haplotype of Ayu maintained individually in experimental tanks based on eDNA samples. Second, we examined whether we could correctly detect variation in mitochondrial haplotypes from an eDNA sample containing nine haplotypes derived from 20 individuals of Ayu. We prepared multiple PCR replicates for an eDNA sample and analyzed them separately. We compared the number of true haplotypes and false positive haplotypes between the results obtained with and without the use of two major ASV methods (DADA2 and UNOISE3) for processing the HTS data. During the analysis, we put special emphasis on the detection rate of each haplotype in PCR replicates because erroneous sequences are expected to occur randomly during experimental processes (e.g., PCR and MiSeq sequencing), and false positive haplotypes are therefore expected to rarely be detected in multiple PCR replicates. Here we expected that false positive haplotypes could be eliminated correctly from HTS data of an eDNA sample by using ASV methods and/or removing haplotypes with low detection rates among PCR replicates.

## Material and Methods

### Ethics statement

The fish used for our experiments were purchased from a local fisherman. Based on current laws and guidelines of Japan relating to animal experiments on fish, the collection of fish tissue for extracting DNA and the use of DNA samples are allowed without any ethical approvals from any authorities. However, all experiments were performed with attention to animal welfare.

### Primer design

We targeted the mitochondrial D-loop region, because it has a higher mutation rate compared with the nuclear DNA regions and the other mtDNA regions (Moritz et al. 1987). To amplify the control region of Ayu, two sets of species-specific primers were developed based on the complete mitochondrial DNA sequence of Ayu from the National Center for Biotechnology Information (NCBI; http://www.ncbi.nlm.nih.gov/, accession numbers of downloaded sequences were AB047553, EU124679–EU124683). The first primer set, Paa-Dlp-1 primer, was developed for Sanger sequencing, which can amplify nearly the entire D-loop region (amplicon length, 541 bp). The sequences of the primers are as follows:

PaaDlp-1_F

(5 ‘-GCTCCGGTTGCATATATGGACC-3’),

PaaDlp-1_R

(5 ‘-AGGTCCAGTTCAACCTTCAGACA-3’)

The second primer set, PaaDlp-2, was designed for HTS by referring to instructions suggested previously (Miya et al. 2015; Palumbi 1996). We obtained 232 sequences of the mtDNA control region from Ayu from the MitoFish v.2.80 (Iwasaki et al. 2013; http://mitofish.aori.u-tokyo.ac.jp/) and aligned these sequences using mafft. The information for all sequence data used to design primers for HTS (PaaDlp-2 primers) is listed in Table S1. The aligned sequences were imported into MESQUITE v. 2.75 (Maddison et al. 2010), and the search for a short hypervariable region (up to 200 bp for paired-end 150 bp sequencing using the Illumina MiSeq) flanked by two conservative regions (ca 20–30 bp) was performed in the entire region of aligned sequences. For HTS, we designed the PaaDlp-2 primers to be within the amplification range of PaaDlp-1, considering the unconventional base pairing in the T/G bond to enhance the primer annealing (i.e. the designed primers use G rather than A when the template is variable C or T, and T rather than C when the template is A or G, Fig. 1). Two types of reverse primers, PaaDlp2_1R and PaaDlp2_2R, were designed, because the reverse priming site has one variable nucleotide (the template is A or G) that does not bind despite the T/G bond. The base R indicates A (PaaDlp-2_R1) or G (PaaDlp-2_R2). The primer sequences are as follows:

**Figure 1.**
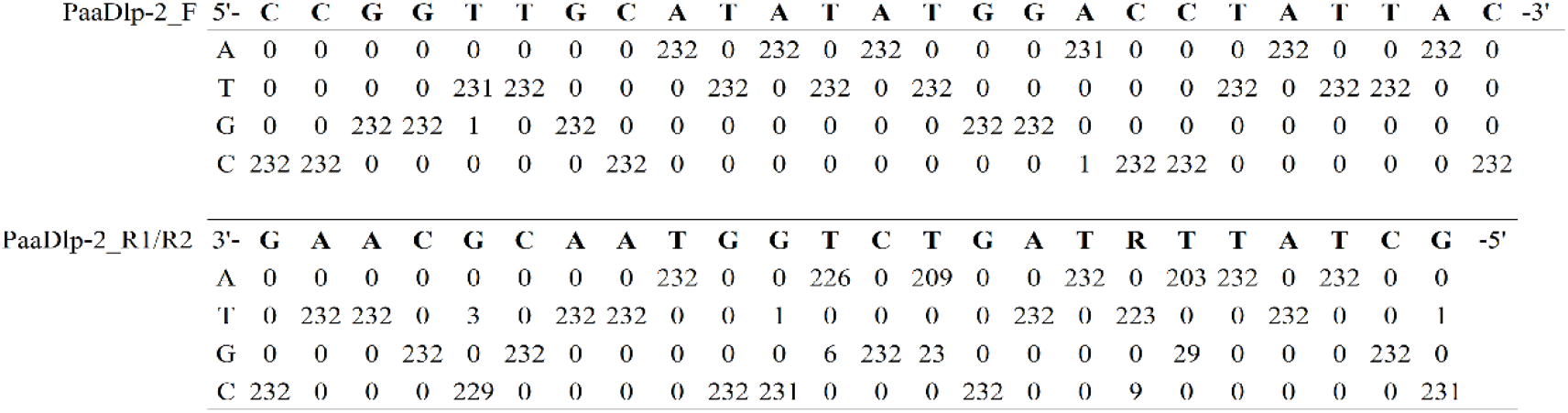
The sequences of the PaaDlp-2 primers and sequence variation in the corresponding region (Downloaded data from MitoFish v.2.80.). Two types of reverse primers, PaaDlp2_1R and PaaDlp2_2R, were designed, because the reverse priming site has one variable nucleotide (the template is A or G) that does not bind despite the T/G bond. Note the presence of nucleotide substitutions only in one sequence out of 232, which was ignored during primer design.

PaaDlp-2_F

(5’-CCGGTTGCATATATGGACCTATTAC−3’),

PaaDlp-2_R1 and PaaDlp-2_R2

(5’-GCTATTRTAGTCTGGTAACGCAAG−3’).

To check the specificity of the PaaDlp-1(F/R) and PaaDlp-2(F/R1/R2) primers, we performed an *in silico* specificity screen using Primer-BLAST with default settings (http://www.ncbi.nlm.nih.gov/tools/primer-blast/).

### Haplotype determination from tank water eDNA and corresponding individuals

We obtained 20 juveniles of Ayu (0.92 ± 0.21 g wet weight, mean ± SD) that were caught with a large fixed net in Lake Biwa (35°18’25” N; 136°3’40” E, DMS) in Japan on 24 February 2015. Live fishes were brought back to the laboratory and then maintained individually in a small tank with 300 mL of aged tap water at room temperature. After 15 min, each fish was removed from the tanks and anaesthetized with an overdose of clove oil. To extract DNA from the tissues, about 0.02 g of skeletal muscle tissues was collected from each individual. DNeasy Blood & Tissue Kit (Qiagen, Hilden, Germany) was used following the manufacturer’s protocol.

To collect eDNA, 250 mL of rearing water was sampled from each tank and vacuum-filtered using a Whatman GF/F glass fiber filter (GE Healthcare Japan, Tokyo, Japan; diameter 47 mm; nominal pore size of 0.7 μm). All filter disks were folded in half inward with tweezers and wrapped in aluminum foil, then stored at −20°C. eDNA was extracted according to the methods described in the section, ‘ eDNA extraction from filters’. As a filtration negative control (FNC), the same volume of ultrapure water was filtered in the same manner after the filtration of the all the real samples. The FNC was treated alongside the samples in the following experimental steps to confirm no cross contamination. Before use, all sampling and filtration equipment were exposed to a 10% bleach solution for 10 min, washed with running tap water and rinsed with ultrapure water. PCR was performed in a 25-μL reaction for each sample using the StepOnePlus Real-Time PCR System. The mixture of the reaction was as follows: 900 nM each of PaaDlp-1(F/R) in 1 × PCR master mix (TaqMan gene Expression Master Mix, Life Technologies, Carlsbad, CA, USA) with 7 μL of sample eDNA or 1 μL of tissue-derived DNA (10 ng/μL). The PCR thermal conditions were 2 min at 50°C, 10 min at 95°C, and 44 cycles of 15 s at 95°C and 60 s at 60°C. The PCR products were purified using Nucleo Spin^®^ Gel and PCR Clean-up Kits (Code No. 740609.50; TAKARA Bio, Kusatsu, Japan) according to the manufacturer’s instructions. Sequences were determined by commercial Sanger sequencing service (Takara Bio, Kusatsu, Japan). The sequences which were successfully determined (total 448 bp) were deposited in the DNA database of Japan (DDBJ, https://www.ddbj.nig.ac.jp/dra/index.html; accession numbers, LC406364- LC406383) and are listed in Table S2.

### eDNA extraction from filters

The filter samples were subjected to eDNA extraction following the method described in Yamanaka et al. (2017). The filter was rolled into a cylindrical shape using sterile forceps and placed in the upper part of a spin column (EZ-10; Bio Basic, Markham, Ontario, Canada) from which the silica membrane had been removed before use. Excess water remaining in the filters was removed by centrifugation for 1 min at 6000 *g*, and a mixture of 200 μL of ultrapure water, 100 μL of Buffer AL and 10 μL of proteinase K was dispensed onto the filter in each spin column and incubated for 15 min at 56°C. The Buffer AL and proteinase K were supplied from the DNeasy Blood & Tissue Kit. After incubation, the spin columns were centrifuged for 1 min at 6000 *g* to elute the eDNA into 2-mL tube. The upper part of the spin column was placed in a 2-mL tube, and 200 μL of Tris-EDTA buffer (pH 8.0) was added to the filter and incubated for 1 min at room temperature to recover the remaining DNA on the filter. The spin columns were centrifuged for 1 min at 6000 *g* to obtain the second elution and mixed with the first elution. Subsequently, 100 μL Buffer AL and 600 μL ethanol were added to each tube and mixed by pipetting. The DNA contained in mixture was purified using a DNeasy Blood and Tissue Kit following the manufacturer’s protocol and eluted using 100 μl of the elution buffer provided with the kit.

### Detection of mitochondrial haplotype diversity from an eDNA sample

The experimental design is shown in Fig. 2. We collected 50 mL of rearing water from each tank used in the experiment described above. All collected water was mixed (total volume 1 L) and vacuum-filtered using one Whatman GF/F glass fiber filter. Environmental DNA was extracted according to the methods described above. After extracting eDNA from the filter sample, we employed a two-step tailed PCR approach to construct paired-end sequencing libraries, according to methods described by Miya et al. (2015). The FNC sample in the first experiment was used again as FNC sample in this experiment, because the filtrations of both experiments were performed at the same time. To avoid the risk of cross-contamination, all sampling and filtering equipment were decontaminated with 10% bleach solution for more than 10 min, carefully washed with tap water, and finally rinsed with ultrapure water. In addition, to safeguard eDNA samples against cross-contamination, filtered pipette tips were used and separate rooms were used for pre- and post-PCR operations.

**Figure 2.**
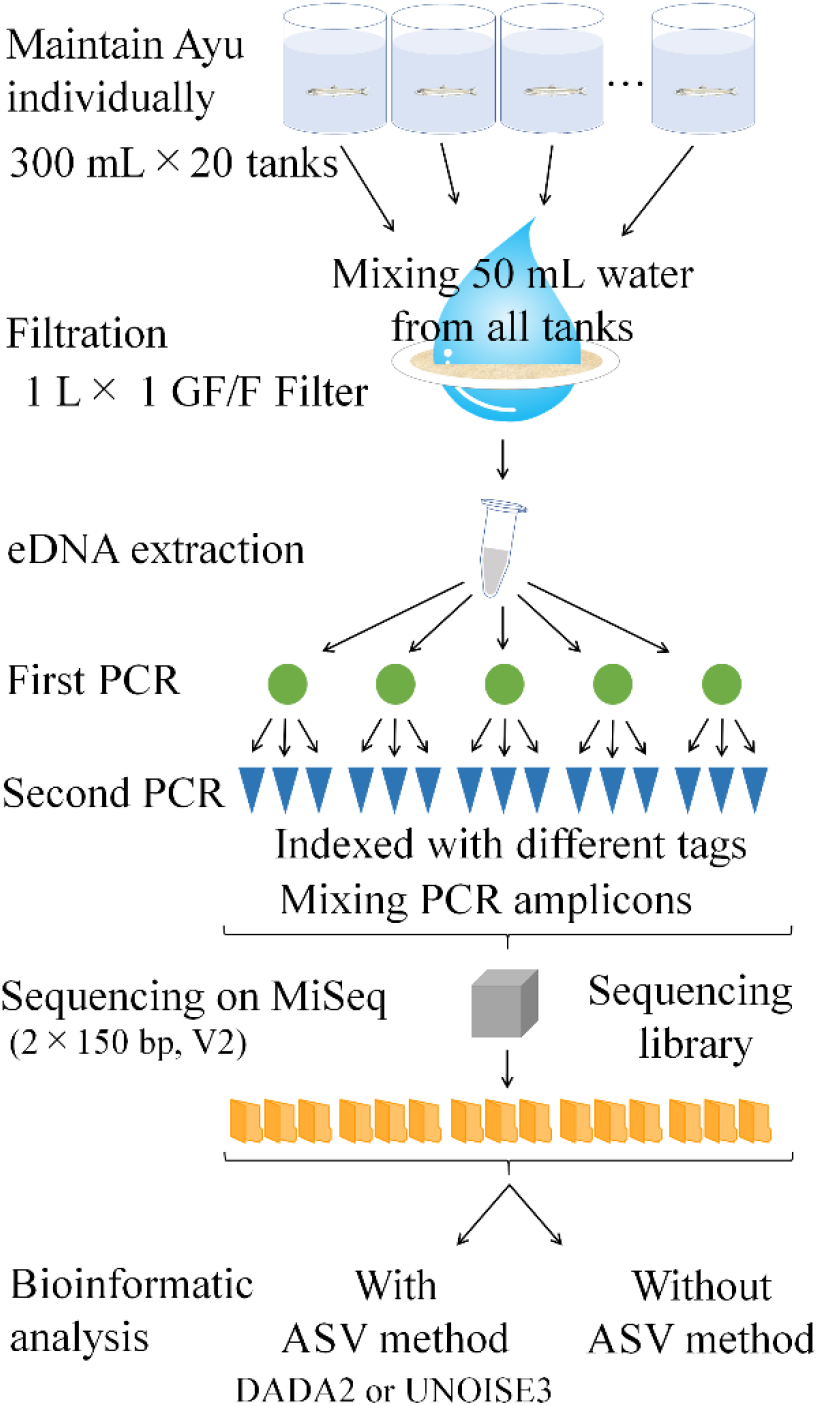
Experimental design for detecting mitochondrial haplotype diversity from an eDNA sample.

The first PCR was performed in five replicates, each in a 12-μL reaction. The target region of Ayu was amplified using primers containing adapter sequences and random hexamers (N). The primer sequences are as follows: 5’-ACACTCTTTCCCTACACGACGCTCTTCCGATCTNNNNNN + PaaDlp-2_F (gene-specific sequences) −3’ and 5’-GTGACTGGAGTTCAGACGTGTGCTCTTCCGATCTNNNNNN + PaaDlp-2_R1/R2 (gene-specific sequences) −3’. The mixture of the reaction was as follows: 0.3 μM of PaaDlp-2_F, 0.15 μM each of PaaDlp-2_R1 and R2 in 1 × KAPA HiFi HotStart ReadyMix (KAPA Biosystems, Wilmington, WA, USA) and a 2-μL sample of eDNA. To monitor cross contamination during library preparation, nontemplate control (NTC) were included in triplicate in the first PCR. The PCR thermal conditions were 3 min at 95°C, 35 cycles of 20 s at 98°C, 15 s at 60°C, and 15 s at 72°C, followed by a final extension for 5 min at 72°C. The first PCR products were purified twice using Agencourt AMPure XP beads (Beckman Coulter), according to the manufacturer’s instructions (reaction ratio; AMPure beads 0.8: PCR product 1, target amplicon length; ca. 290 bp).

The second PCR was performed in three replicates, with each a 12-μL reaction of the first PCR (total 15 replicates). To distinguish PCR replicates during Illumina MiSeq sequencing, respective PCR replicates (total 15 replicates) were indexed with different combinations of indexing primers. The primer sequences used in second PCR are listed in Table S3. The mixture of the reaction was as follows: 0.3 μM of each second PCR primer in 1 × KAPA HiFi HotStart ReadyMix and 2 μL of the purified first PCR product from the Agencourt AMPure XP beads. As negative controls in the second PCR, 2 μL of the first PCR product of the NTC was added to each reaction instead of template eDNA. The PCR thermal conditions were 3 min at 95°C, 12 cycles of 20 s at 98°C and 15 s at 65°C, with a final extension for 5 min at 72°C. The indexed second PCR products were pooled in equal volumes (5 μL each). The target size of the libraries (ca. 370 bp) was collected using 2% E-Gel^®^ SizeSelect™ agarose gels (Thermo Fisher Scientific, Waltham, MA, USA) according to the manufacturer’s instructions.

DNA concentrations in the collected libraries were estimated using the Qubit fluorometer (Life Technologies) with the Qubit dsDNA HS assay kit and adjusted to 4 nM (assuming 1 bp equals 660 g mol^−1^) using ultrapure water. A 5-μL volume of the 4 nM library was denatured with 5 μL of fresh 0.2 N NaOH, followed by 5 μL of Tris HCl (200 mM, pH 7) and 985 μL of HT1 buffer (including in Miseq Reagent Kit) was added to adjust the library concentration to 20 pM. Then, 48 μL of 20 pM PhiX DNA (Illumina, San Diego) and 360 μL of HT1 buffer were added to 192 μL of the 20 pM library to obtain an 8 pM library. The library was sequenced using the MiSeq platform (Illumina, San Diego), with the MiSeq v2 Reagent Kit for the 2 × 150 bp PE cartridge (Illumina, San Diego). The sequencing reads obtained in the present study were deposited in the DDBJ Sequence Read Archive (accession number: DRA006638).

The MiSeq paired-end sequencing (2 × 150 bp) of the 21 libraries for this study (containing 15 PCR replicates, three FNC and three NTC), together with 105 libraries from another study (total number of libraries =126), yielded a total of 15.98 million reads, with 97.5% base calls containing Phred quality scores greater than or equal to 30.0 (Q30; error rate = 0.1% or base call accuracy = 99.9%).

In this experiment, ‘unexpected’ haplotypes such as heteroplasmy, somatic mutation, and pseudo-gene etc. are expected to detect from HTS data. Because these “unexpected” haplotypes are infrequently present in all organisms, they are not usually detected by direct Sanger sequencing. Therefore, we postulate that a haplotype obtained by Sanger sequencing of tissues samples represents the true haplotype of each individual.

### Bioinformatic analysis

The full range of amplicons obtained using the PaaDlp-2 primers (166bp) were successfully sequenced using the MiSeq platform. However, for the amplicons obtained using the PaaDlp-1 primers, some bases following the forward primer were undetermined by Sanger sequencing of the tissue-derived DNA from 20 individuals of Ayu. The forward primers of PaaDlp-2 and PaaDlp-1 were designed to be close to each other, and thus, three bases after the forward primer of PaaDlp-2 needed to be omitted to compare the overlapping regions between the two datasets. Therefore, only 163 of the bases successfully determined for the two datasets were used for the subsequent bioinformatic analyses. To perform a denoising for erroneous sequences based on the ASV methods, fastq files containing raw reads were processed using the Divisive Amplicon Denoising Algorithm 2 package ver. 1. 12. 1 (DADA2, Callahan et al. 2016) of R and the UNOISE3 (Edgar 2016a). The core algorithm of DADA2 infers unique biological variants using denoising algorithm that is based on a model of errors in the amplicon sequencing with MiSeq. The detailed algorithm of DADA2 is described in the original paper (Callahan et al. 2016). Briefly, the adopted error model in DADA2 quantifies the rate *λji*, at which an amplicon read with sequence *i* is produced from sample sequence *j* as a function of sequence composition and quality. Then, the *p*-value of the null hypothesis that the number of amplicon reads of sequence *i* is consistent with that of the error model is calculated using a Poisson model for the number of repeated observations of the sequence *i*, parameterized by the rate *λji*. Calculated p-values are used as a division criterion for an iterative partitioning algorithm, and sequence reads are divided until all partitions are consistent with being produced from their central sequence. Reads of sequences inferred as errors are replaced with the central sequence of the partition that included its sequence (i.e. error correction). Quality-filtered sequences are dereplicated, and the parameters of the DADA2 error model are trained on a random subset of one million reads. The trained error model is used to identify and correct indel-mutations and substitutions. Denoised forward and reverse reads are merged and read pairs with one or more conflicting bases between the forward and reverse read are removed. In this study, trimming of primer and random hexamers were performed using the function ‘removePrimers’, and then, reads with one or more expected errors (maxEE = 1) were discarded during quality inspection. DADA2 implements the function ‘ removeBimeraDenovo’ to identify chimeras; however, it was not used in this study because haplotypes of Ayu included in the sampled water might be incorrectly identified as chimeras due to high sequence similarity.

To predict whether a sequence is correct or not, the UNOISE3 computes the probability of incorrectness of a sequence based on the abundance skew ratio (ratio of abundance) and number of differences between sequences. The detailed algorithm of UNOISE3 is described in the original paper (Edgar 2016a). Briefly, the adopted error model in UNOISE3 divides sequences into clusters each of which has a centroid with high-abundance unique read *C*. The cluster includes the sequence *M* with abundance *aM* which has close sequence similarity with the centroid *C*. The number of differences including both substitutions and gaps between *M* and *C* is defined as the Levenshtein distance d. Neighbor sequences of *C* with small *d* and small *aM* are predicted to be error reads of *C*. In this study, the denoising by UNOISE3 was performed using UNOISE3 option implemented in Usearch v11 (https://drive5.com/usearch/manual/). The threshold of the minimum abundance (size= annotation) was set at 4 reads, and length filter was set at 100 bp, respectively. To identify chimeras, the UCHIME2 (Edgar 2016b) implemented in Usearch v11 was used.

Furthermore, the HTS data was also processed without the ASV method. The base calling errors were eliminated by quality filtering. The data preprocessing and dereplicating were performed using a custom pipeline described by Sato et al. (2018). Briefly, the low-quality tails were trimmed from each read, and the tail-trimmed paired-end reads (reads 1 and 2) were assembled using the software FLASH with a minimum overlap of 10 bp. The primer sequences were then removed with a maximum of three mismatches. Only when sequences had 100% identity with each other, they were operationally considered as identical. The fastq format was transformed into fasta, and the pre-processed reads were dereplicated. At this point, the reads were subjected to a local BLASTN search against a custom-made database of the control region of Ayu. The custom-made database was constructed from the 190 haplotypes of mitochondrial D-loop region of Ayu, which were downloaded from MitoFish (Table S4) and were obtained from the 96 individuals caught at Ado river (35°19’31” N, 136°3’48” E, DMS, Japan). The Ado river is connected to Lake Biwa and located close to a large fixed net (ca. 4 km) that was used to catch Ayu juveniles for the present study. The information in all haplotypes included in the custom-made database is listed in Table S4. If the respective sequences obtained in the HTS data had ≥ 99% similarity with the reference haplotype and an E-value < 10^−5^ in the BLAST results, the sequences were identified as those of Ayu.

### Statistical analysis

All statistical analyses were performed using R ver. 3. 6. 0 software (R Core Team. 2019) and the minimum level of significance was set at *α* = 0.05. To determine differences in the total number of reads derived from the nine true haplotypes, which were derived from 20 individuals (see “Results”), and false positive haplotypes, a Mann–Whitney *U* test was performed. A generalized linear model (GLM) with Poisson distribution was used to test how well the detection rate of false positive haplotypes in the PCR replicates correlated with the total reads of the false haplotypes (glm function in R ver. 3.6.0 software).

## Results

### Testing species specificity of the two primer sets

The PaaDlp-1_F/R and PaaDlp-2_F/R1/R2 had mismatches within five bases from the 3’ end of them with closely related species *(Hypomesus nipponensis*) which is sympatrically distributed with Ayu (Fig. S1). The *in silico* specificity check for PaaDlp-1 and PaaDlp-2 (no adapter sequence) implemented in Primer-BLAST indicated species-specific amplification of Ayu. The direct sequencing of the PCR amplicons corroborated the amplification of the target region of Ayu in section ‘Haplotype determination from tank water eDNA and corresponding individual’.

### Comparison of detected haplotypes from tissue-derived DNA and corresponding tank eDNA

The sequences from the 20 Ayu individuals that were used for tank experiments were classified into 17 and nine haplotypes based on Sanger sequencing of PCR products amplified using PaaDlp-1 (amplicon length: 448 bp) and PaaDlp-2 (amplicon length: 166 bp), respectively (Table S2 and Fig. S2). The detected haplotypes had only one or a few differences from each other, with the maximum pairwise *p*-distances for the two datasets being 0.022 (PaaDlp-1) and 0.025 (PaaDlp-2), respectively. Each sequence obtained from eDNA, which was amplified using PaaDlp-1 and was detected from each of the 20 rearing water tanks, was identical to that obtained from tissue-derived DNA of the corresponding individual. In this tank experiment, the target fragments were not detected in any FNC and NTC. Thus, there was no evidence for cross-contamination during sample processing.

### Detection of mitochondrial haplotype diversity from an eDNA sample

There were no non-target species sequences in the all sequence data obtained by two types of ASV methods. Based on the bioinformatic analysis using the DADA2, 1,639,277 reads were detected and assigned to 46 haplotypes (Fig. 3A, Table S5). Of these, 1,555,097 (94.9%) reads were assigned to nine true haplotypes and they were detected from 15/15 of PCR replicates. The remaining 84,137 (5.1%) reads were assigned to 37 false positive haplotypes. A total of 34 (0.002%) and 43 (0.003%) reads were detected from the three FNC and three NTC (Table S5), respectively. The 37 false positive haplotypes were detected at rates ranging between1/15 to 15/15 PCR replicates, but nine false positive haplotypes were detected in all 15 PCR replicates. In addition, the false positive haplotypes with a low detection rate were randomly detected from the 15 PCR replicates and were not derived from any particular first PCR replication (Table S5).

**Figure 3.**
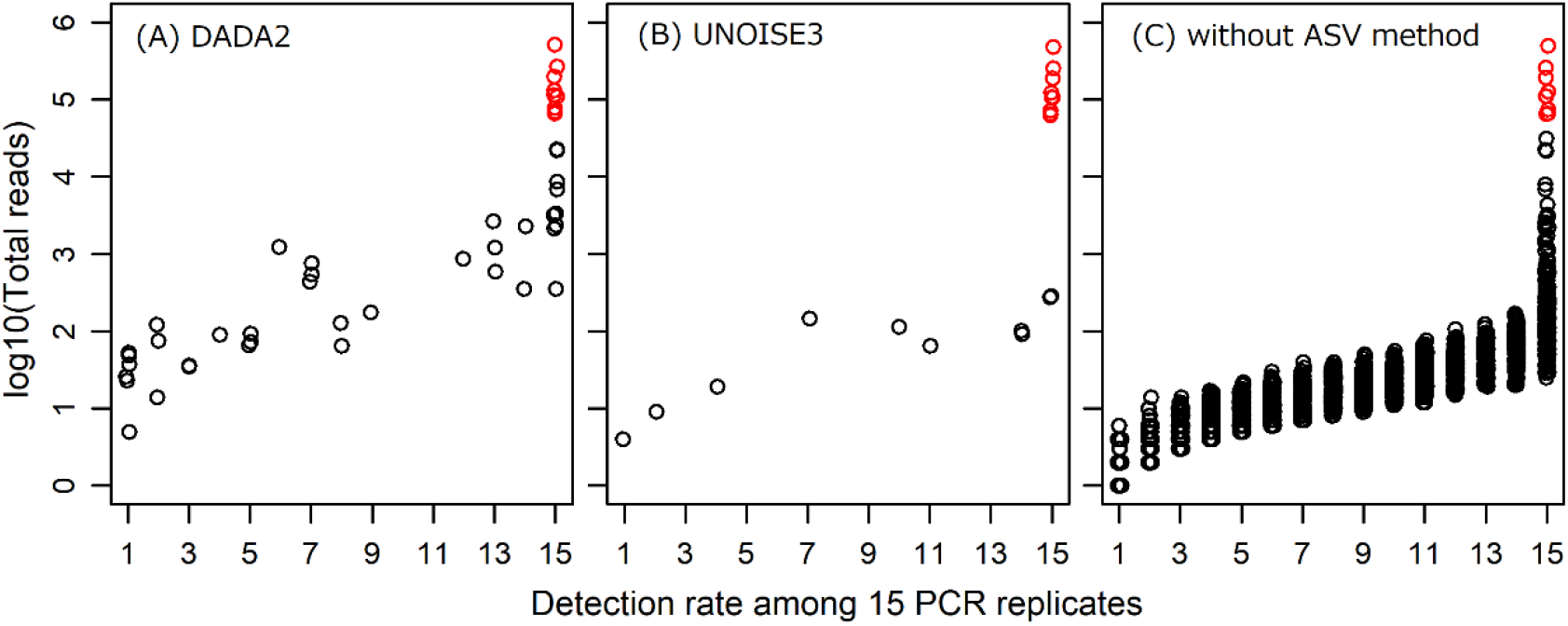
Relationships between detection rate and total reads for each haplotype. They were detected with (A) DADA2, (B) UNOISE3, and (C) without the ASV method. The red and black circle indicate the true haplotype and false positive haplotype, respectively.

Based on the bioinformatic analysis using the UNOISE3, 1,466,578 reads were detected and assigned to 19 haplotypes (Fig. 3B, Table S5). Of these, 1,465,402 (99.9%) reads were assigned to nine true haplotypes and they were detected from 15/15 of PCR replicates. The remaining 1,113 (0.1%) reads were assigned to 10 false positive haplotypes. A total of 32 (0.002%) and 31 (0.002%) reads were detected from the three FNC and three NTC (Table S5), respectively. The 10 false positive haplotypes were detected at rates ranging between1/15 to 15/15 PCR replicates, but two false positive haplotypes were detected in all 15 PCR replicates. In addition, the false positive haplotypes with a low detection rate were randomly detected from the 15 PCR replicates and were not derived from any particular first PCR replication (Table S5).

Based on the bioinformatic analysis without the ASV method, 1,748,030 reads out of the total reads that passed quality control processes were assigned to Ayu with greater than or equal to 99% identity to the reference haplotypes in the custom-made database. Of these, 1,502,828 (86%) reads were assigned to nine true haplotypes, and they were detected from 15/15 PCR replicates (Fig. 3C). The remaining 245,202 (14%) reads consisted of 5,683 false positive haplotypes. The 5,683 false positive haplotypes were detected in 1/15 to 15/15 detection rates; however, 335 false positive haplotypes were detected in 15/15 PCR replicates (Fig. 3C). Despite the efforts to decrease the risk of cross-contamination, 124 (0.007%) and 105 (0.006%) reads were detected from the three FNC and three NTC, respectively. Regardless of whether the ASV methods was used, read abundances of the true haplotypes were significantly larger than those of the false positive haplotypes (Mann-Whitney *U* test; DADA2, *p*< 0.001; UNOISE3, *p*< 0.001; without ASV method, *p* < 0.001; Fig. 3). Furthermore, the total read count of false positives increased significantly with increasing detection rate (GLM;*p* < 0.001, *p* < 0.001,*p* < 0.001; Fig. 3).

## Discussion

We found that correcting or eliminating erroneous sequences with two types of ASV methods, DADA2 and UNOISE3, was effective to improve the accuracy of intraspecific genetic diversity evaluations with eDNA analysis. Furthermore, the accuracy of the analysis seems to be further improved by removing of haplotypes with low detection rates. Although some caution is still required for risk of false positives and false negatives, the proposed approach is useful for applying eDNA analysis to evaluation of intraspecific genetic diversity that requires higher accuracy with respect to distinguishing true haplotypes from false ones.

The use of the both ASV methods in eDNA analysis for evaluating intraspecific genetic diversity considerably decreased the number of false positive haplotypes (Fig. 3). In addition, both of ASV methods detected all the true haplotype with 15/15 detection rate. The great performance of the ASV methods for eliminating false positive haplotypes is consistent with previous studies that identified microorganisms from mock community samples at fine taxonomical resolutions (Callahan et al. 2016; Hughes et al. 2017; Kopylova et al. 2016). In previous studies, the detection of true/false haplotypes relied on known reference sequences (Sigsgaard et al. 2016, Parsons et al. 2018), while the ASV-based approach used in the present study allowed us to distinguish true haplotypes from false positive haplotypes without the reference database. This is especially advantageous when a target species does not have sufficient reference sequences. Therefore, the use of ASV methods will expand the applicability of eDNA-based evaluation of intraspecific genetic diversity for various species. However, it still needs further studies in outdoor to make a robust conclusion on which ASV method (DADA2, UNOISE3, or others) is more suitable for eDNA-based evaluation of genetic diversity.

The present results suggest that detection rate of each haplotype in PCR replicates provides an important clue to discriminate true haplotypes from false positive haplotypes. The accuracy of eDNA-based evaluation of genetic diversity would be further increased by selecting only haplotypes with high detection rates among multiple PCR replicates. True haplotypes, especially predominant haplotypes, would be amplified at an early stage of PCR in all PCR replicates. Thus, detection rates of true haplotypes in PCR replicates are expected to be much higher than those of false positive haplotypes, which was clearly supported in this study (Fig. 3). On the other hand, false positive haplotypes would not be contained in the initial eDNA template, but it was considered that they were incidentally generated during the first and second PCR and/or sequencing step of HTS (see Fukui et al. 2013; Nakamura et al. 2011). To identify the step at which the false positive haplotypes were introduced, we tried to follow-up the detection pattern based on separate tags in each PCR replicate. However, there was no particular detection pattern among 15 PCR replicates, and it was difficult to identify the step at which most of false positive haplotypes from our data generated.

The present study also has implications for understanding the relationship between the total reads of each haplotype and the number of individuals owning that haplotype. Previous studies have suggested that the eDNA concentration increases with an increase in abundance and/or biomass of organisms (Doi et al. 2016; Pilliod et al. 2013; Takahara et al. 2012; Yamamoto et al. 2016). In addition, eDNA sequence reads are likely to be correlated with biomass and the number of individuals belonging to each fish family in a community (Thomsen et al. 2016). Thus, there is a possibility that the total number of reads of each true haplotype reflects the number of individuals that represent corresponding haplotypes (hereafter called ‘owner individuals’). In this study, using the GLM analysis with the Poisson distribution, the number of owner individuals had a significant positive effect on the total reads of the haplotypes regardless of whether the ASV methods was used (*p* < 0.01 in all cases; Fig. 5). This result suggests that the use of eDNA analysis has the potential to evaluate not only the diversity of haplotypes but also the relative dominance of each haplotype in a population. In addition, this result is consistent with that in Sigsgaard et al. (2016) that showed a correlation between the relative abundances of respective haplotypes in a population and those in eDNA samples. Furthermore, the total read abundance potentially can be used to eliminate false positive haplotypes based on an appropriate threshold value. However, we consciously avoided this approach. In a field setting, *a priori* information on the haplotype composition in the target population and the relative concentration of eDNA corresponding to each haplotype are usually unavailable, and thus, the use of a higher threshold value for the total read abundance may lead to eliminate true haplotype and increase false negative rates (Elbrecht et al. 2018).

**Figure 4.**
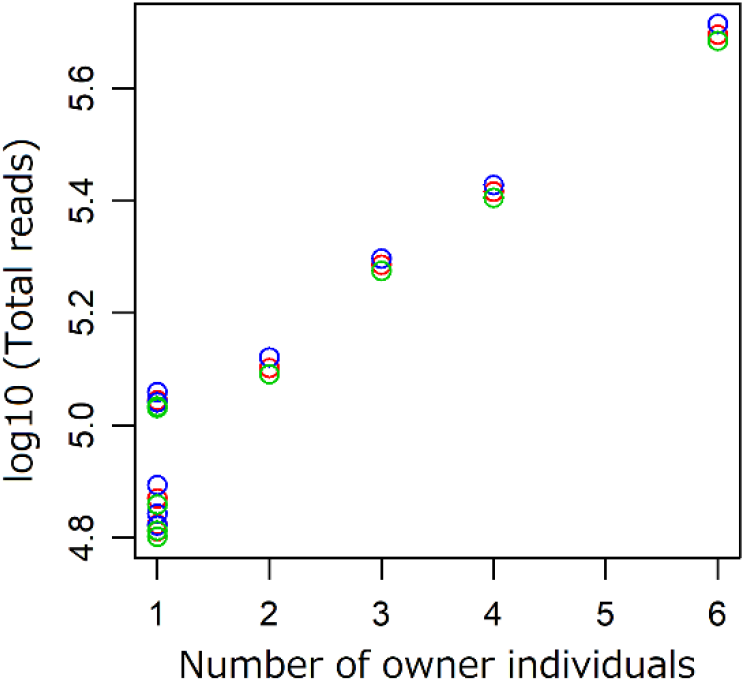
Relationship between the number of owner individuals and total reads for each detected true haplotype. Blue, green, and red circles indicate true haplotypes detected with DADA2, UNOISE3, and without the ASV method, respectively. The number of owner individuals of each haplotype had a significant positive effect on the total reads in all analysis methods (p < 0.01, p < 0.01, and p < 0.01, respectively).

In future studies, it will be necessary to further test how accurately and comprehensively we can detect true haplotypes in the field, where eDNA concentrations should be lower than those in tank experiments (e.g. Minamoto et al. 2017; Takahara et al. 2012). In addition, heterogeneous distributions of eDNA in field water has been also reported (Dejean et al. 2012; Jerde et al. 2011; Pilliod et al. 2013; Thomsen et al. 2012; Yamamoto et al. 2016). Therefore, it is necessary to determine optimal sampling strategies, containing appropriate volumes of sample water and distances between sampling points to accurately evaluate intraspecific genetic diversity using eDNA analysis. Additionally, the use of protein coding marker (such as Cyt*b* and COI) may have potential to increase the detection accuracy by the denoising. In the protein coding marker, synonymous substitutions at the third codon position usually predominate. Thus, the use of protein coding marker may have potential to distinguish true haplotype from false positive haplotypes based on whether base substitution is synonymous or non-synonymous substitution. Further development of eDNA analysis to evaluate intraspecific genetic diversity would contribute to more effective genetic resource management and ecosystem monitoring.

## Author contributions

i. the conception or design of the study: S.T., T.M., and H.Y
ii. the primer development: S.T. (PaaDlp-1), and M.M (PaaDlp-2).
iii. the acquisition, analysis, or interpretation of the data: S.T., M.M., M.U., and H.S.
iv. writing of the manuscript.: S.T., M.M., M.U., H.S., T.M., and H.Y.

## Data Archiving Statement

The minimal raw dataset is uploaded to the DDBJ Sequence Read Archive (https://www.ddbj.nig.ac.jp/dra/index-e.html; Accession number: DRA006638).

## Acknowledgements

We thank Mr. Sakurai S. and Mr. Shibata N. (Ryukoku University) for supporting the experiments. We thank Dr. Yamamoto S. (Kyoto University), Dr. Tanabe A.S. (Ryukoku University), Dr. Fukaya K. (National Institute for Environmental Studies), and Doi H. (Hyogo University) for constructive comments and suggestions to this study. We thank Dr. Callahan B.J. for supporting the analysis in DADA2. His advises and kind supports have helped us to improve the manuscript. This work was supported by JSPS KAKENHI Grant Number JP16K18610 and Grant-in-Aid for JSPS Research Fellow Grant Number JP18J10088.

**Supporting Information 1**

Table S1

Title: The information of all sequence data which were used for designing the PaaDlp-2 primers.

**Supporting Information 2**

Table S2

Title: Detected sequence haplotypes from 20 individuals of Ayu.

**Supporting Information 3**

Table S3

Title: Primer sequences for second PCR.

**Supporting Information 4**

Table S4

Title: The information of all haplotypes included in custom-made database.

**Supporting Information 5**

Table S5

Title: Haplotypes detected with the ASV methods and the read counts for each haplotype in each PCR replicate.

**Supporting Information 6**

Fig. S1. Alignment of priming sites of Ayu *(Plecoglossus altivelis altivelis)* and its closely related species *(Hypomesus nipponensis*) which sympatrically distributed. F1 and R1 indicate the priming sequence of PaaDlp-1_F and PaaDlp-1_R, respectively. F2 and R2 indicate the priming sequence of PaaDlp-2_F and PaaDlp-2_R1/R2, respectively. Asterisks indicate Ayu-specific nucleotides.

**Supporting Information 7**

Fig. S2

Title: Neighbour-joining tree of detected haplotypes from 20 Ayu individuals using (a) PaaDlp-1 and (b) PaaDlp-2.

