## Supplementary material for "Evaluating intraspecific genetic diversity of a fish population using environmental DNA: An approach to distinguish true haplotypes from erroneous sequences": Fig. S1

**Supporting Information 6**

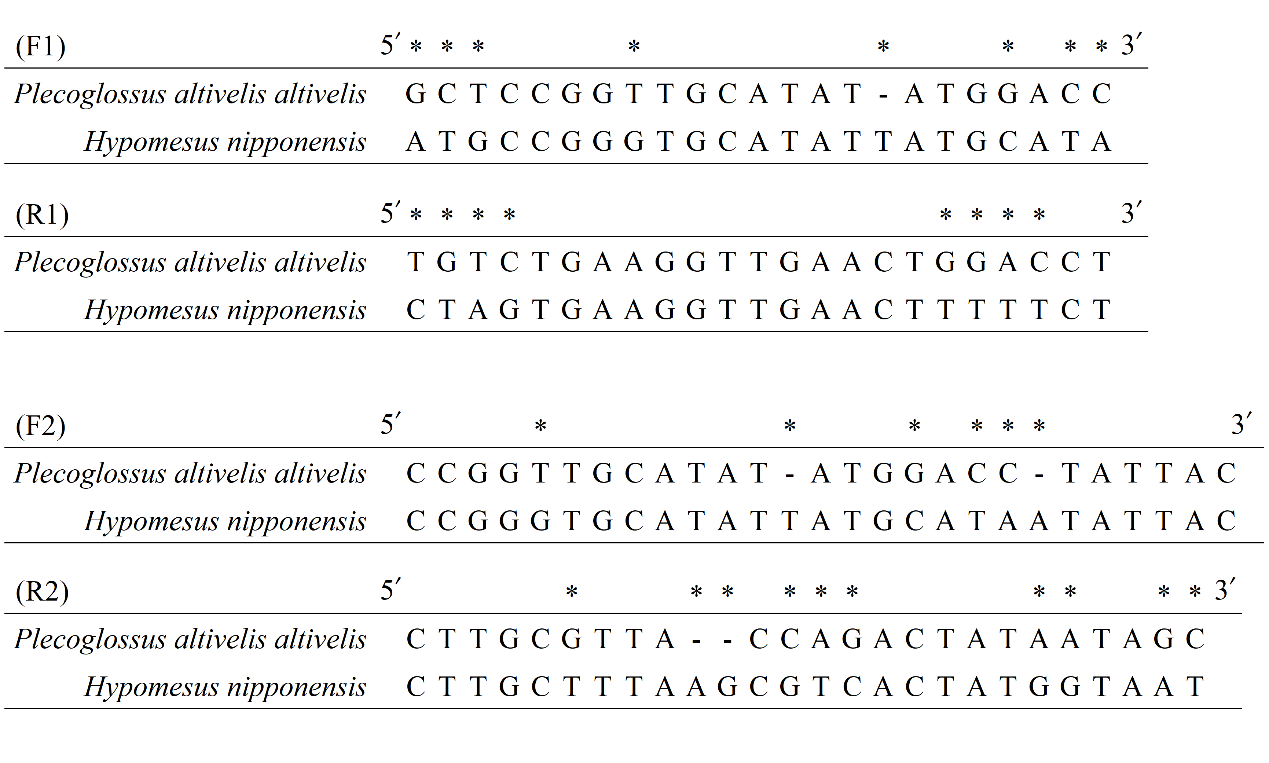

Fig. S1

Alignment of priming sites of Ayu (*Plecoglossus altivelis altivelis*) and its closely related species (*Hypomesus nipponensis*) which sympatrically distributed. F1 and R1 indicate the priming sequence of PaaDlp-1_F and PaaDlp-1_R, respectively. F2 and R2 indicate the priming sequence of PaaDlp-2_F and PaaDlp-2_R1/R2, respectively. Asterisks indicate Ayu-specific nucleotides.
