## Supplementary material for "Evaluating intraspecific genetic diversity of a fish population using environmental DNA: An approach to distinguish true haplotypes from erroneous sequences": Fig. S2

**Supporting Information 7**

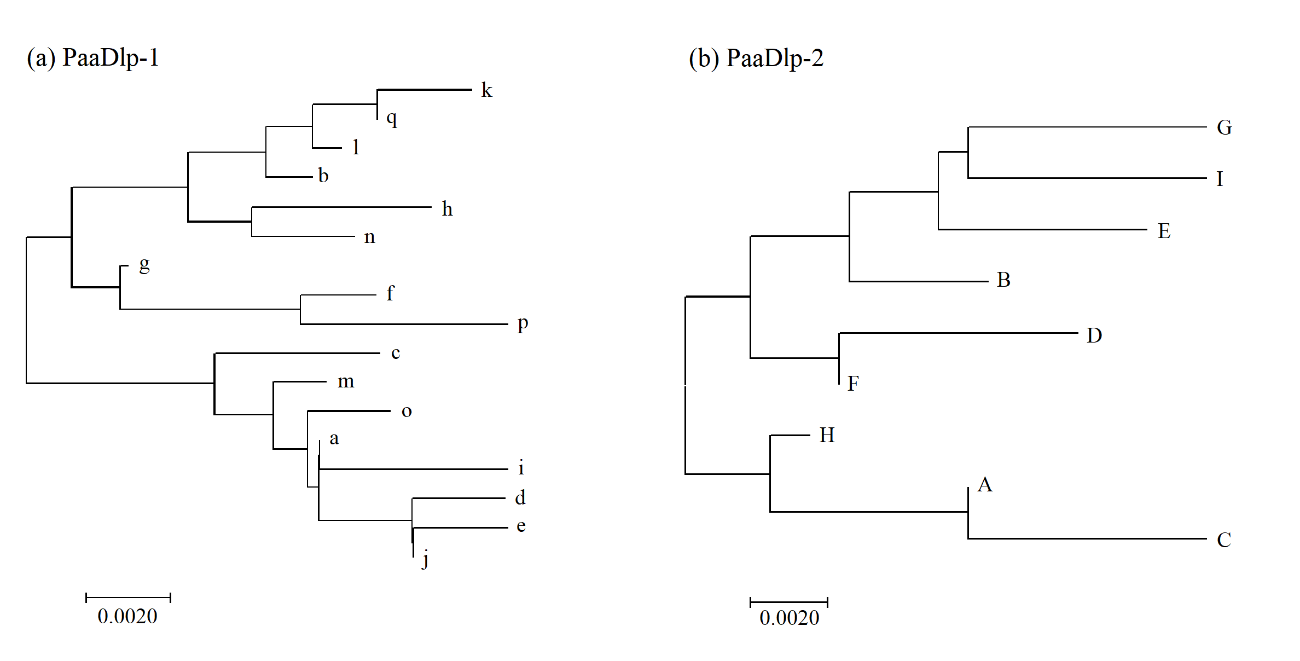

Fig. S2

**Neighbour-joining tree of detected haplotypes from 20 Ayu individuals using (a) PaaDlp-1 and (b) PaaDlp-2.** The tree was constructed using the p-distance method in MEGA ver. 6.0.6 software (Tamura et al., 2013). The scale bar indicates the number of base differences per site as the evolutionary distances between haplotypes.
